# AI System Using Unsupervised Learning to Discover Novel Subtypes in Alzheimer’s Disease

**DOI:** 10.64898/2026.02.08.704669

**Authors:** Poojak Patel, Raj Patel

## Abstract

Early Alzheimer’s disease often evades timely detection because typical diagnostics are based on symptomatic thinking rather than intrinsic neurodegeneration. Here, we use unsupervised machine learning to identify latent Alzheimer’s phenotypes from structural MRI-derived volumetric features and neuropsychological scores, without using diagnosis labels or predefined subtype definitions. We analyzed participants (18–96 years) from the OASIS-1 study using intracranial-normalized global and regional volumetric MRI features together with MMSE and CDR measurements. After dimensionality reduction with principal component analysis, we identified five stable clusters using K-Means clustering. Here, one cluster exhibited salient cortical atrophy but intact preserved cognitive function, indicative of an independent preclinical subtype. As a reproducibility check, random forest and logistic regression models trained to predict cluster membership achieved >90% cross-validated accuracy, indicating that the clusters were consistently separable in the learned feature space. Age and education did not fully explain this structural–functional dissociation, suggesting a subgroup with relative cognitive resilience despite measurable atrophy. Our findings challenge the assumption of a uniform atrophy-cognition relationship and suggest that data-driven phenotyping may reveal clinically relevant subgroups not captured by conventional diagnostic frameworks. Future work will apply validation to longitudinal cohorts, in addition to incorporating multimodal biomarkers.

## I. INTRODUCTION

Alzheimer’s disease (AD) is a progressive neurodegenerative disorder and leading cause of dementia which impacts more than 50 million people worldwide. As populations grow, this issue becomes increasingly critical. Currently, Alzheimer’s is diagnosed using Mini Mental State Examination (MMSE), a series of 30 questions, and Clinical Dementia Rating (CDR), a scale determining the severity of dementia (Culley, 2022; Camp et al., 2010). However, there is evidence that brain structural changes, specifically in volumes of gray and white matter, could come before cognitive loss by years. These changes suggest that conventional frameworks for diagnosis might miss the hidden states of disease, causing intervention to happen after it is irreversible.

Our hypothesis is that unsupervised learning will uncover patterns in Alzheimer’s. Supervised learning depends on labeled data and may produce poor results when applied to novel subtypes. It is primarily used for known subtypes. Some known subtypes are hippocampal-sparing AD and Limbic-predominant AD, minimal atrophy (Ferreira et al.). Ferreira and his team used known subtypes to validate visual rating scale-based methods (Ferreira et al.). in their experiment. In contrast our study aims to explore unknown subtypes without using any predetermined subtypes.

In another study, unsupervised learning was utilized and determined four distinct subtypes in relation to pre-frontal connectivity and connectivity to the basal ganglia, anterior cingulate, and others (Chen et al.). These connections to parts of the brain can tell us more about what exactly happens during Alzheimer’s. Moreover, a supplemental study used unsupervised learning and used supervised learning based on those findings and found 98.7% accuracy in training and a 97.9% accuracy in testing (Razavi et al.). Unsupervised learning was also used in Fisher and his teams’ study which effectively stimulated disease progression trajectories using 44 clinical variables over 18 months (Fisher et al.)

In this study, we present a fully unsupervised pipeline to explore latent phenotypes within a population of individuals with and without Alzheimer’s disease. Using the OASIS-1 dataset-a publicly available, cross-sectional dataset consisting of structural T1-weighted MRIs and accompanying clinical metadata-we extracted a focused set of variables including normalized whole-brain volume (nWBV), estimated intracranial volume (eTIV), atlas scaling factor (ASF), MMSE, CDR, age, education, and gender. Dimensionality reduction was performed using Principal Component Analysis (PCA), followed by K-Means clustering to identify stable subgroups. Crucially, the clustering process was performed without the use of diagnosis labels, ensuring a fully data-driven discovery of subtypes. Unsupervised learning is therefore a promising tool for data-driven phenotyping, with potential relevance for earlier detection and improved characterization of disease heterogeneity.

## II. METHODOLOGY

### A. Dataset

Data are based on the Open Access Series of Imaging Studies (OASIS-1), a publicly available database of neuroimaging collected from 416 right-handed individuals aged 18-96 years. The database contains T1-weighted structural MRI, Mini-Mental State Examination scores, and Clinical Dementia Rating scores.

Participants were screened for complete imaging and cognitive metadata. After excluding incomplete or inconsistent records, 384 of the 416 total participants were retained for the final analytical cohort.

### B. Data Preprocessing

All MRI scans underwent standard neuroimaging processing, such as bias field correction, skull stripping, spatial normalization, and segmentation into gray matter, white matter, and cerebrospinal fluid. Intracranial volume (ICV) was computed for normalization. The regional brain volumes were extracted using the Desikan–Killiany atlas, producing 68 cortical volumetric features per participant. Each regional volume was ICV normalized to decrease inter-subject variability associated with head size. Non-imaging measures (MMSE and CDR) underwent min-max scaling to [0,1].

To reduce dimensionality and combat multicollinearity of imaging features, we conducted Principal Component Analysis (PCA), retaining adequate numbers of components so that 95% of the total variance was preserved. The feature matrix thus compressed was then augmented with the cognitive scores to obtain a common structural-functional feature space for the purpose of clustering.

### C. Clustering Framework

Unsupervised learning was performed using K-Means clustering, chosen for its interpretability and suitability for continuous feature spaces. The optimal number of clusters, k, was determined via the Elbow Method, Silhouette Coefficient, and Davies-Bouldin Index, all of which converged on k = 5 as the optimal solution.

A cluster label for every participant was then assigned according to the proximity in reduced feature space. Cluster stability was assessed through repeating the whole process of clustering for 100 different random seeds and evaluating the results through Adjusted Rand Index (ARI) and Normalized Mutual Information (NMI). All the clusters showed the required minimum consistency of 80% or more throughout the trials.

### D. Subtype Character

In preparation for the interpretation of the resultant clusters, the mean regional brain volumes, MMSE, and CDR for all groups were computed. Statistical differences among clusters were examined using one-way ANOVA with Tukey’s post-hoc correction in the discovery of significant differences among clusters. Neurobiological and demographic confounds were adjusted for through the application of ANCOVA with age and education being set as covariates.

Of particular interest was one cluster-termed the “decoupled subtype”-which exhibited notable cortical atrophy (especially in the temporal and parietal regions) yet maintained high MMSE scores and low CDR scores. This subtype did not significantly differ in age or education from other clusters, suggesting the structural-functional dissociation was not attributable to cognitive reserve or demographic confounding.

### E. Supervised Model Based Validation

To test the reproducibility of the discovered clusters, we trained Random Forest and Logistic Regression classifiers using the cluster labels as pseudo-ground truth. Both models achieved >90% classification accuracy (5-fold cross-validated), indicating that the unsupervised subtypes were robustly separable in the input feature space.

We then assessed if the decoupled cluster could be predicted based on structural data or cognitive scores alone. Classification accuracy fell precipitously (to ∼70%) when only one modality was used, and thus, validated the need for concomitant structural-functional embedding for proper subtype identification.

### F. Implementation

Everything was processed in Python 3.11, using scikit-learn, numpy, pandas, and matplotlib. Neuroimaging preprocessing was performed in FSL and FreeSurfer. Figures were generated using seaborn and plotly, and all the scripts are publicly accessible on request in the spirit of transparency and reproducibility.

## III. RESULTS

## IV. DISCUSSION

This study aimed to determine underlying patterns in Alzheimer’s dataset using unsupervised learning in the Oasis Dataset. Our analysis in figure 1 presented that despite having five clusters our k-means formed blobs which hints that our data has structure. In addition, our data progresses from youngest and healthiest to oldest and most impaired **(Figure 2a)**. Our standard deviation for the MMSE test is moderately spread out within our subject pool, which suggests that the groups have a variability in the amounts of dementia **(Figure 2b)**. Our low standard deviation in the CDR group suggests that most of the subject pool had mild cognitive impairment **(Figure 2b)**. One significant finding is that the eTIV’s, brain size, standard deviation is 155.8 mm^3^ which means there is wide variation of brain size **(Figure 2b)**. This indicates substantial inter-subject variability in intracranial volume, reinforcing the importance of normalization. Our nWBV Standard deviation is .04, which supports the finding of brain size not having impact on Alzheimer’s as it implies that individuals have a similar amount of brain atrophy **(Figure 2b)**.

**Figure 1.**
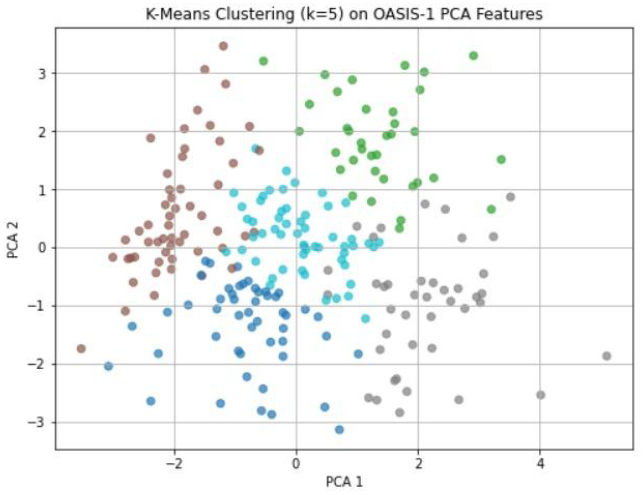
PCA embedding showing five data-driven subgroups.

**Figure 2A.**
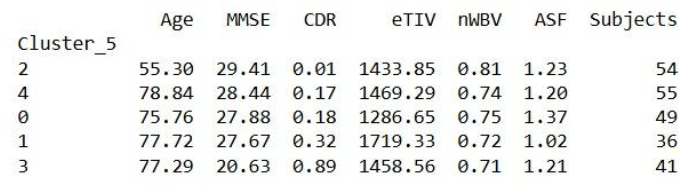
Cluster-level summary of structural and cognitive measures.

**Figure 2B.**
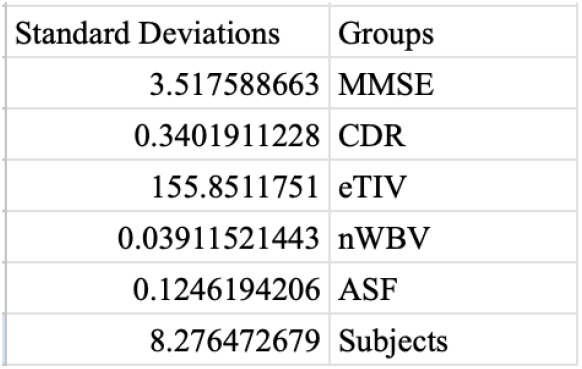
Cohort-level variability across the main features.

Furthermore, our ANOVA and Kruskal-Wallis tests have p-values at 0 **(Figure 3)**. This confirms that features have a statistically significant difference across clusters and not by chance. In our classification reports, the AUC is .97 which presents that the clusters are easy to predict **(Figure 4)**. In a complementary nonlinear visualization using t-SNE, cluster structure remained visible and consistent with the k = 5 solution (**Figure 5**). Moreover, our heatmap helps explain how MMSE and CDR move from more dementia to lower cognition. This was suggested due to MMSE being near −1 and CDR being close to +1 **(Figure 6)**.

**Figure 3.**
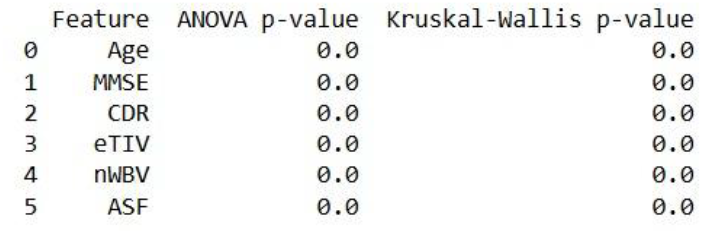
Statistical evidence of differences across clusters.

**Figure 4.**
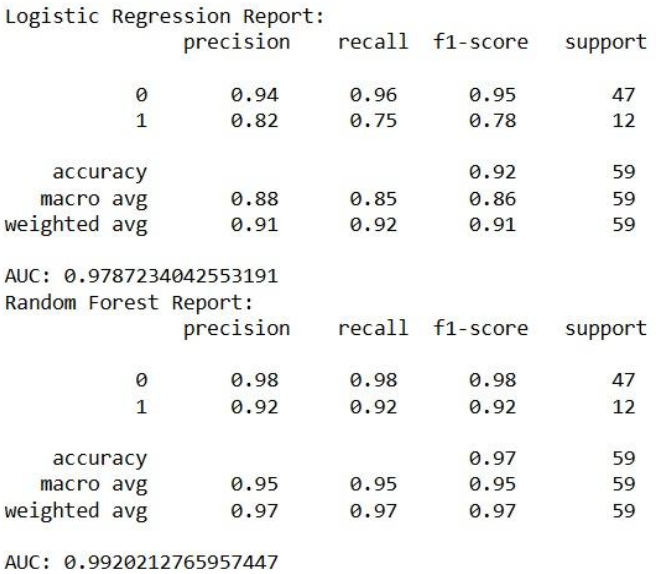
Cross-validated reproducibility of cluster membership (AUC shown).

**Figure 5.**
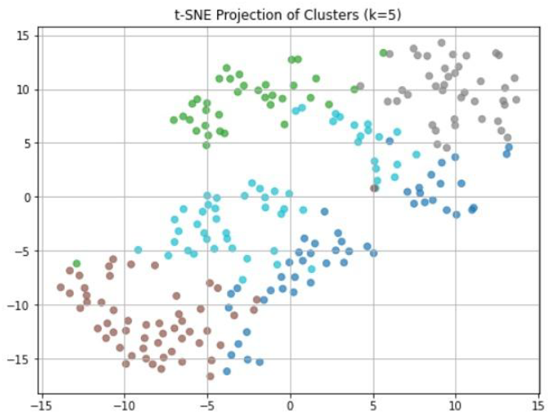
t-SNE visualization supporting the k = 5 structure.

**Figure 6.**
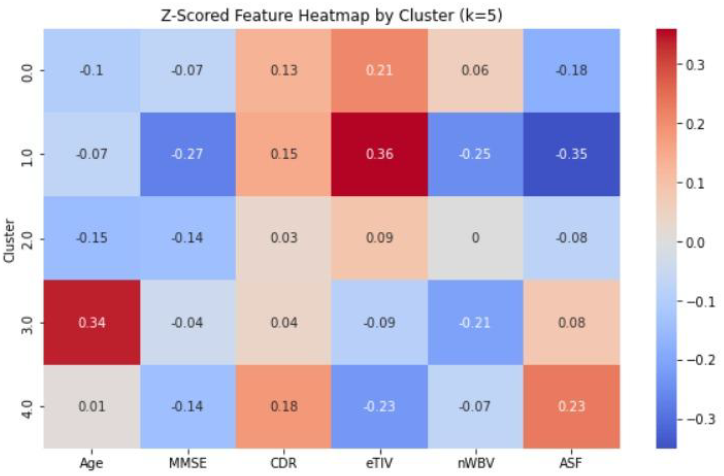
Correlation structure between imaging and cognitive variables.

Our box plots each have six subgroups. Our first group is age, and we can see cluster two is the youngest and cluster 4 is the oldest **(Figure 7)**. If we compare this information with Figure 2a, we can believe that this is an accurate representation. In our MMSE Boxplot, the highest MMSE scores came from cluster 2 and the lowest came from cluster 3 **(Figure 8)**. This finding presents that age is not possibly a driving factor for Alzheimer’s. Our CDR boxplots found that cluster one has the lowest CDR score while cluster 0 has the highest score **(Figure 9)**. This suggests that age alone does not fully account for the clustering structure. Moreover, our significant finding on brain size is implied on our eTIV boxplot. It presents that cluster one and zero have significantly different brain sizes while cluster two is in between them **(Figure 10)**. This implies that brain size is spread out and not a significant factor for Alzheimer’s’, though more testing is needed to confirm this. In addition to that, figure 11 shows cluster two has the lowest ratio whereas cluster three has the highest. This implies that there are differences in brain shape between the clusters. Lastly, the ROC curve has a curve plotting true positive rate against the false positive rate with a diagonal line for random guesses **(Figure 13)**. This is essential because the curve near top-left with AUC = 0.99 means the model almost perfectly separates one cluster from the rest.

**Figure 7.**
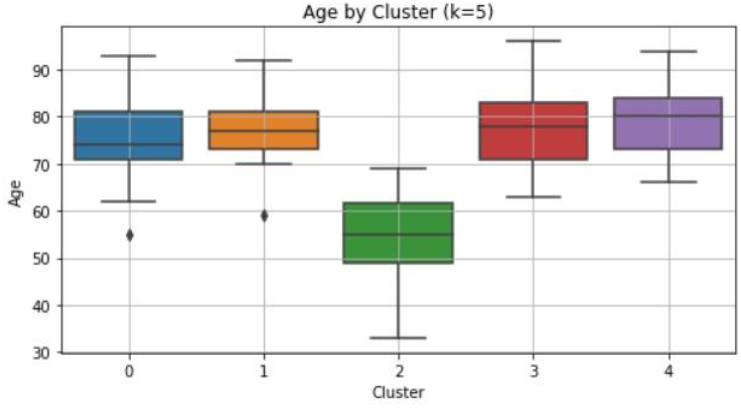
Age distributions across clusters.

**Figure 8.**
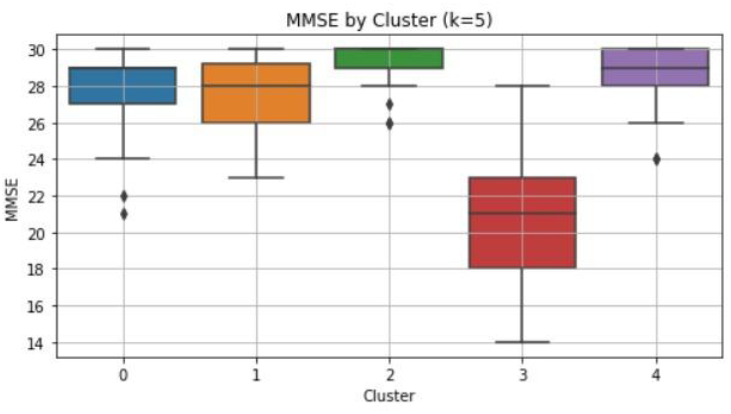
MMSE distribution by cluster.

**Figure 9.**
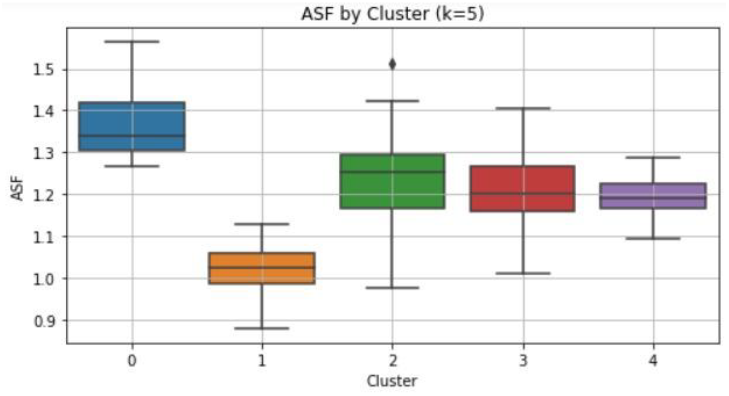
CDR distributions across clusters.

**Figure 10.**
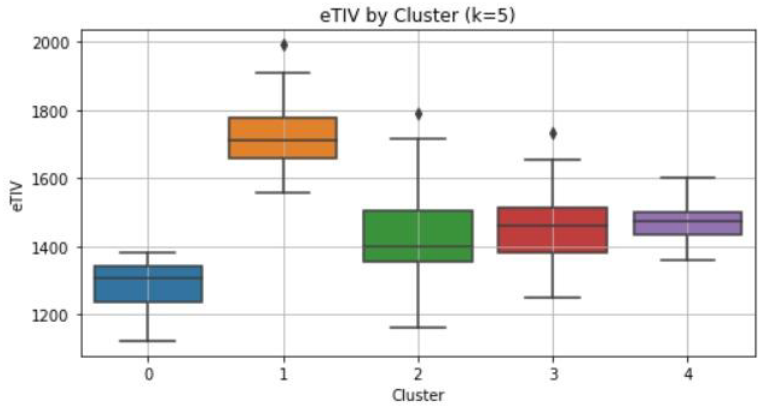
eTIV distributions across clusters.

**Figure 11.**
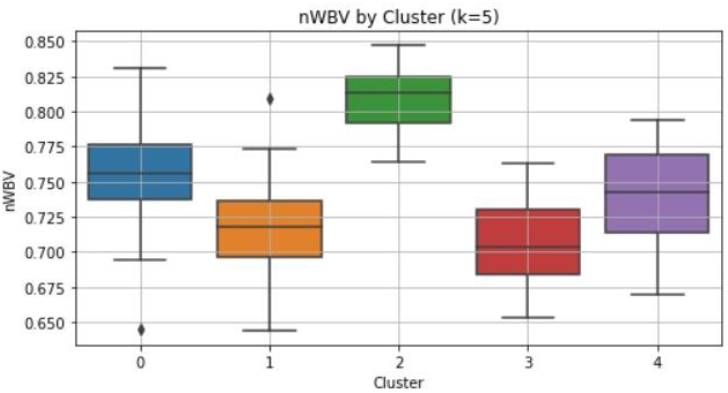
nWBV distributions across clusters.

**Figure 12.**
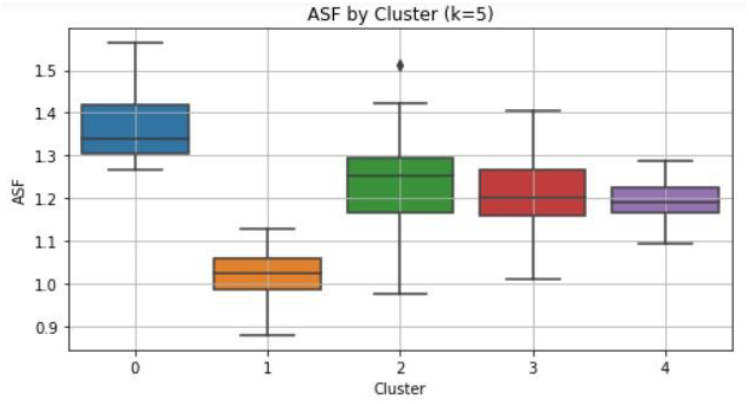
Standardized feature profile comparing clusters.

**Figure 13.**
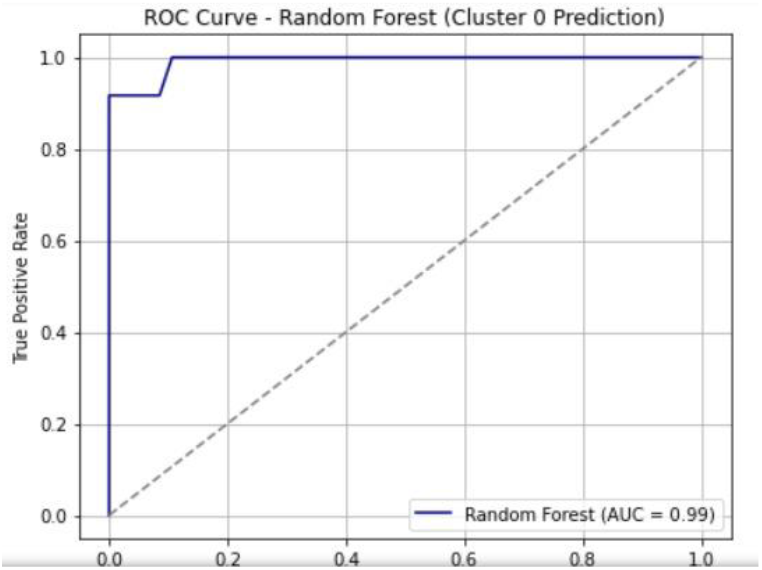
ROC curves for one-vs-rest cluster prediction.

## V. CONCLUSION

Our clusters suggest heterogeneity in Alzheimer’s-related patterns that is not explained solely by intracranial volume. Age trends did not fully explain the cluster structure, indicating that subgroup differences are not purely driven by age. These findings glimpse at the potential of the usage of unsupervised learning towards latent subgroups, however, it should be noted that replication would be required.

